# The cap-independent translational regulation of *NbRabGAP1* in *Nicotiana benthamiana* during BaMV infection

**DOI:** 10.64898/2026.01.07.698292

**Authors:** Jia-Ling Shih, I-Hsuan Chen, Ying-Ping Huang, Ying-Wen Huang, Ching-Hsiu Tsai

## Abstract

While cap-dependent translation is the eukaryotic norm, alternative pathways enable selective protein synthesis during stress. We describe a cap-independent mechanism regulating NbRabGAP1, a host factor essential for bamboo mosaic virus (BaMV) movement in *Nicotiana benthamiana*. The 5′ untranslated region (5′ UTR, 553-nt) of *NbRabGAP1* contains three upstream open reading frames (uORFs) that typically hinder ribosome scanning. Using bicistronic reporter assays, we identified a functional internal ribosome entry site (IRES) within nucleotides 61-150. This IRES features a predicted stable Y-shaped RNA structure and a purine-rich R-motif. We demonstrate that NbPAB8, a poly(A)-binding protein, specifically binds this R-motif to enhance IRES-mediated translation over 2.5-fold. During BaMV infection, *NbPAB8* expression is transiently upregulated, coinciding with maintained *NbRabGAP1* levels despite potential repression of global cap-dependent translation. This represents a regulatory switch where viral infection triggers a PAB8-dependent IRES mechanism to sustain the vesicle trafficking machinery required for viral movement. Our findings reveal how viral stress rewires host translational control through specific RNA-protein interactions, ensuring the synthesis of critical host factors via IRES-mediated pathways.

**Highlight:** During bamboo mosaic virus infection, NbPAB8 binds a purine-rich IRES in the NbRabGAP1 5′ UTR, bypassing global translation repression to maintain vesicle trafficking essential for viral movement.

## Introduction

In eukaryotes, protein translation occurs through two fundamental mechanisms: cap-dependent and cap-independent translation. The predominant cap-dependent translation pathway relies on recognition of the 5′ cap structure by the translation initiation complex, which scans the mRNA from the 5′ end toward the 3′ direction until encountering the first AUG initiation codon (Dever, 1999; Kozak, 1978). Subsequently, the 60S ribosomal subunit joins the 48S preinitiation complex to form the 80S ribosome and initiate translation (Kozak, 1992). Under certain condition, however, such as pathogen infection or heat stress, ribosomes can bypass the cap structure and directly bind to the 5′ untranslated region (UTR) at a specific element called an internal ribosome entry site (IRES). IRES elements form distinct secondary structures that recruit initiation factors and facilitate direct association of the 40S ribosomal subunit (Kozak, 2001; Pestova *et al*., 1996).

In animals, the concept of IRES-mediated translation was first discovered in picornaviruses, where ribosomes initiate translation internally within the 5’ UTR (Pelletier and Sonenberg, 1988). During virus infection, host cap-dependent translation is often suppressed either by cleavage of cap-binding protein by viral protease (Imataka and Sonenberg, 1997; Lamphear *et al*., 1995; Mader *et al*., 1995; Sachs and Varani, 2000) or by competition for translation factors due to massive viral RNA accumulation (Lopez-Ulloa *et al*., 2022). It is therefore conceivable that certain cellular mRNAs also contain IRES elements in their 5′ UTRs, enabling their translation when cap-dependent initiation is compromised. The highly structured nature of many 5′ UTRs can hinder ribosome scanning, further favoring cap-independent initiation. Indeed, IRES-mediated translation of host mRNAs has been documented under various physiological stresses, such as pro-survival stress adaptation or cell death (Komar and Hatzoglou, 2011).

Compared with their viral counterparts, cellular IRES elements are structurally more diverse and often less stable (Xia and Holcik, 2009). They function through complex interactions with canonical initiation factors, IRES *trans*-acting factors (ITAFs), and the 40S ribosomal subunit (Komar and Hatzoglou, 2011). In the plant kindom, the 5′ UTR of Hsp101 in *Zea mays* contains a stem-loop-based IRES element that recruits the eIFiso4G initiation factor to promote cap-independent translation during heat stress (Dinkova *et al*., 2005; Jimenez-Gonzalez *et al*., 2014). Similarly, the 5′ UTR of WUSCHEL in *Arabidopsis thaliana* mediates IRES-driven translation in shoot apical meristem cells in response to environmental hazards (Cui *et al*., 2015).

Rab GTPases are small GTP-binding proteins that play essential roles in intracellular vesicle trafficking among eukaryotic organelles. They regulate vesicle budding, cytoskeletal transport, targeting, docking, and membrane fusion, in cooperation with multiple accessory factors for activity and recycling (Grosshans *et al*., 2006; Stein *et al*., 2003). Rab GTPase-activating proteins (RabGAPs) stimulate the intrinsic GTPase activity of Rab proteins, hydrolyzing GTP to GDP and facilitating vesicle transit from donor to acceptor compartment (Novick and Zerial, 1997; Zerial and McBride, 2001). Intracellular vesicle trafficking is also implicated in viral transport (Carluccio *et al*., 2014; Genoves *et al*., 2010; Helenius, 2018; Serra-Soriano *et al*., 2014). In *Nicotiana benthamiana*, Rab proteins such as NbRabF1 and NbRabGAP1 have been shown to participate in bamboo mosaic virus (BaMV) infectioin, facilitating vesicle trafficking that support viral cell-to-cell movement (Carluccio *et al*., 2014; Huang *et al*., 2013; Huang *et al*., 2020; Huang *et al*., 2016; Xu and Nagy, 2016).

In our previous work, we cloned and sequenced the full-length cDNA of *NbRabGAP1* (Huang *et al*., 2013). Subsequent analysis revealed that its 5′ UTR is unusually long (553 nucleotides) and contains three upstream open reading frames (uORFs). In this study, we aim to elucidate the role of this 5′ UTR in the translational regulation of *NbRabGAP1*, particularly in the context of BaMV infection.

## Materials and methods

### Constructs

To investigate the potential IRES function of the 5′ UTR of *NbRabGAP1*, we utilized the bicistronic vector p2Luc (Grentzmann *et al*., 1998) for *in vitro* translation. The known IRES in the 5′ UTR of classical swine fever virus (CSFV) (Huang *et al*., 2012a) was amplified by PCR using the primer set, 5′CSFV-IRES (5′-<u>GTCGAC</u>GTATACGAGGTTAGTTCATT-3′; *Sal*I site underlined) and 3′CSFV-IRES (5′-<u>GGATCC</u>GTGCCATGTACAGCAGAGAT-3′; *Bam*HI site underlined). The fragments of the 5′ UTR of *NbRabGAP1* and the 5′ UTR with removal of the initiation codon (NbRabGAP1 5′UTR/ΔAUG) were PCR amplified with the primer sets, 5′GAP1 (5′-<u>GGATCC</u>TTAAGACCATACAAAAACAAAC-3′, *Bam*HI site underlined)/ 3′GAP1 (5′-<u>GAGCTC</u>CATTTTTTGACCAAAATTGTA-3′, *Sac*I site underlined); and 5′GAP1 and 3′GAP1/ΔAUG (5′-<u>GAGCTC</u>TTTTTGACCAAAATTGTAAT-3′, *Sac*I site underlined), respectively. These fragments were cloned into p2Luc via restriction enzyme sites, *Bam*HI and *Sac*I.

To modify the vector p2Luc by inserting a stem-loop, we used the primer set 5′p2Luc/SL (5′-G<u>TCTAGA</u>GATCCAGATCCCACCACGGCCCAGATCTGGGCCGT-3′, *Xba*I site underlined) and 3′p2Luc/SL (5′-GGGTACC<u>GGATCC</u>AGATCCCACCACGGC-3′, *Bam*HI site underlined) to anneal each other. The resulting fragment was cloned into p2Luc via *Xba*I and *Bam*HI sites, resulting in a new plasmid vector denoted as p2Luc/SL. Fragments of various lengths of the 5′ UTR of *NbRabGAP1* were PCR amplified using the GAP1 clone as a template and cloned into p2Luc/SL. The full-length of the 5′ UTR (GAP1/1-553) was amplified with primer set, 5′GAP1/1 (5′-G<u>ACTAGT</u>CATATGGACCATACAAAAACAAAC-3′, *Spe*I site underlined) and 3′GAP1/553 (5′-<u>GAGCTC</u>CATTTTTTGACCAAAATTGTA-3′, *Sac*I site underlined). GAP1/1 and GAP1/1-110 were amplified with the primer sets 5′GAP1/1 and 3′GAP1/322 (5′-<u>GAGCTC</u>CATAAGAAAATAGCTGAAATTAAAG-3′, *Sac*I site underlined) or 3′GAP1/110 (5′-<u>GAGCTC</u>CATTTTTGATTTACCCTTTTC-3′, *Sac*I site underlined), respectively. GAP1/322-553 was amplified using the primer set 5′GAP1/322 (5′-G<u>ACTAGT</u>CATATGTTGTAATTTTTTCCCGTTAA-3′, *Spe*I site underlined) and 3′GAP1/553. GAP1/345-480 was amplified with the primer set 5′GAP1/354 (5′-G<u>ACTAG</u>TCATATGGAATCTTGGCTCTAATTTT-3′, *Spe*I site underlined) and 3′ GAP1/480 (5′-<u>GAGCTC</u>CATGAATCAATCAACCCAACAACCAG-3′, *Sac*I site underlined). GAP1/61-150, GAP1/61-240, and GAP1/61-322 were using the primer sets, 5′GAP1/61 (5′-G<u>ACTAGT</u>CATATGTCTACAAAAAGAAAAAACCG-3′, *Spe*I site underlined) and 3′GAP1/150 (5′-<u>GAGCTC</u>CATAGTGGTCTAATCTTGAAGAAT-3′, *Sac*I site underlined), 3′GAP1/240 (5′-<u>GAGCTC</u>CATAAAATAGAGAGAATTGAGCTAAAT-3′, *Sac*I site underlined), or 3′GAP1/322, respectively. As a positive control, the known IRES sequence of maize heat shock Hsp101 was amplified from maize genomic DNA using the primer set 5′Hsp101 (5′-G<u>ACTAGT</u>CATATGTCTAGACTCCCGGCGAAC-3′, *Spe*I site underlined) and 3′Hsp101 (5′-<u>GAGCTC</u>CATGGCTGCTTCTCGGTCCTCAG-3′, *Sac*I site underlined). All these PCR-amplified fragments were cloned into p2Luc/SL via restriction sites *Spe*I and *Sac*I.

To generate the full-length NbPAB8 expression construct, a PCR reaction was performed using BamHI/PAB8F (5′-G<u>GGATCC</u>ATGGCACAGGTTCAGGTGCAGCA-3′, *Bam*HI site underlined) and KpnI/PAB8R/HA (5′-<u>GGTACC</u>TTA*AGCGTAATCTGGAACATCGTATGGGTA*AGAAACCAGACCATC ATTCAG-3′, *Kpn*I site underlined, and HA-tag sequence in italic). The resulting PCR product was then cloned into pUC18 vector. Upon confirmation of the sequence, the resultant clone was digested with *Bam*HI and *Kpn*I. The digested fragment was subsequently subcloned into pEpyon 33k vector

### *In vitro* translation with rabbit reticulocyte lysate (RRL)

The bicistronic vector p2Luc containing a Renilla luciferase (Rluc) gene as the first cistron and a firefly luciferase (Fluc) gene as the downstream cistron. Various intergenic regions were inserted, including the IRES of CSFV (IRES_CSFV_), the reverse sequence of the IRES (IRES_reCSFV_), the 5′ UTR of NbRabGAP1 (NbRabGAP1-5′UTR), and the 5′UTR of NbRabGAP1 without the initiation codon (NbRabGAP1-5′UTR/ΔAUG). The constructs were transcribed and translated *in vitro* using rabbit reticulocyte lysate (TNT RRL) (Promega, Madison, WI, USA; Cat. No. L5020). Each reaction (final volume 12.5 μl) contained TNT RRL, 1 μg DNA plasmid, TNT reaction buffer, T7 polymerase, 1 mM amino acid mixture (minus methionine), and RNase out (5U/μl; Invitrogen, Carlsbad, CA, USA; Cat. No.10-777-019), [^35^S] methionine (>1,000 Ci/mmol at 10 mCi/ml) was added, and the reaction was incubated at 30□ for 3.5 hours. Translation products were separated by 12% SDS-PAGE, and both Rluc and Fluc of each protein fraction were detected.

### *In vitro* transcription and translation

RNA was transcribed from 1 μg *Hin*dIII-linearized plasmid template using T7 polymerase. In each transcription reaction (25 μl), 5 μl of 5X T7 RNA polymerase buffer, 2.5 μl of 25 mM NTP, 2.5 μl of T7 polymerase, and 0.5μl of RNase out (5 U/ μl) (Invitrogen) were combined. The resulting RNAs were quantified and utilized for *in vitro* translation with wheat germ extracts (WGE) (L5030, Promega). Each translation reaction was prepared in a final volume of 10 μl, consisting of 5 μl WGE, 100 ng RNA, 0.4 μl reaction buffer, 0.1 μl 1 mM amino acid mixture (minus leucine), 0.1 μl 1 mM amino acid mixture (minus methionine), and 0.2 μl of RNase out. The reaction was incubated for 1 hour at 30□.

### Dual-luciferase reporter assay

All reagents utilized in the Dual-Luciferase Reporter Assay System (Promega; Cat. No. E1910) were prepared in accordance with the manufacturer’s instructions. Luminescence measurements were carried out following the manufacturer’s recommendations. In a black 96-well microtiter plate, 20 μl of experiment sample was combined with 5 μl of the firefly luciferase reagent (LARII). Luminescence was measured using the SpectraMax L Microplate Reader (Molecular Devices, Silicon Valley, CA, USA). Subsequently, 5 μl of the Renilla luciferase reagent (Stop & Glo) was added, and Rluc luminescence was measured. The data were presented as the ratio of firefly to Renilla luciferase activity (Fluc/Rluc).

### Plant and virus inoculation

*N. benthamiana* plants were cultivated in growth room with a 16/8-hour light/dark cycle at 28 °C. *A. tumefaciens* GV3850 carrying infectious bamboo mosaic virus (pkB) (Liou *et al*., 2014), potato virus X (pkP) (Huang *et al*., 2012b), or an empty vector (pkn) (Prasanth *et al*., 2011) was infiltrated into *N. benthamiana* leaves. To construct the plasmid GAP1/60-GFP, the 5′ UTR of *NbRabGAP1* plus the 60-nts of coding region was amplified with the primer set 5′GAP1+1 (5′-G<u>TCTAGA</u>GACCATACAAAAACAAA-3′, *Xba*I site underlined) and 3′GAP1/+60 (5′-G<u>GGTACC</u>ATCTCCAAAACGACGTGAGGACTCAA-3′, *Kpn*I site underlined). The amplified fragment was then cloned into pBI-mGFP1(GFP only) (Cheng *et al*., 2013) via the *Xba*I and *Kpn*I sites. GFP was fused at the C-terminus of the final product. Plasmids were introduced into *A. tumefaciens* strain C58C1 through electroporation (25 μF, 400 Ω, 2.5□KV pulse), followed by incubation in 1 ml of 2xYT medium at 30 °C with gentle shaking for 2 hours. Transformed cells were plated onto a kanamycin, tetracycline, and rifampicin-containing plate at 30 °C for 2 days. For transient expression in plants, agrobacteria were resuspended in an induction buffer (10 mM MES pH 5.6, 10 mM MgCl_2_, and 150 μM acetosyringone). Co-expression of GAP1-GFP (OD_600_ = 0.4) and GFP (OD_600_ = 0.2) mixed with *Agrobacterium* harboring the silencing suppressor potyviral helper component proteinase (HCPro) (Chiu *et al*., 2010) (OD_600_ = 0.2) and agro-infiltration with pkn (OD_600_ = 0.1) or pkB (OD_600_ = 0.1) was performed. The final density was adjusted to OD_600_ = 0.9 in *N. benthamiana*, and samples were collected at 3 days post-infiltration.

### NbPAB8 expression and purification from *Escherichia coli*

The coding sequence of *NbPAB8* was amplified using the primer set BamHI-PAB8_5 (5′-<u>GGATCC</u>ATGGCACAGGTTCAGGTGCAGCAG-3′, *Bam*HI site underlined) and XhoI-PAB8_3 (5′-G<u>CTCGAG</u>AGAAACCAGACCATCATTCA-3′, *Xho*I site underlined). The PCR product was then cloned into the pGEM-T easy vector (Promega; Cat. No. A1360), and the sequence was verified. Subsequently, NbPAB8 was subcloned from the T-vector into the pET29a expression vector (Addgene, Watertown, MA, USA; Cat. No. 69871-3) and transformed into *E. coli* BL21(DE3).The resulting clone was designated pET29a-NbPAB8. *E. coli* containing pET29a-NbPAB8 was cultured to an OD_600_ of 0.8 in a total volume of 120 ml, and expression was induced with 1 mM isopropyl-D-1-thiogalactopyranoside (IPTG) at 25□ for 16 hr, then samples were centrifuged at 7000 rpm (Hitachi, Himac CR22G) at 4°C for 7 min. The harvested cells were disrupted by sonication in 8 ml of buffer A (20 mM Tris pH 8.0, 100 mM NaCl) containing 10 mM imidazole, protease inhibitor cocktail (Roche, Germany; Cat. No. C755C25) and 1 mM DTT, followed by centrifugation at 12000 rpm (Hitachi, Himac CR22G) at 4°C for 10 min. The supernatant was incubated with 1.5 ml IMAC Ni-Charged Resin (Bio-Rad, Hercules, CA, USA; Cat. No. 780-0801) at 4°C with rotation for 1 hr, then washed with 4.5 ml of buffer A containing 50 mM imidazole, and eluted with buffer A containing 250 mM imidazole. Finally, the eluted protein was dialyzed three times with 350 ml dialysis buffer (20 mM Tris pH 8.0, 100 mM NaCl, 2 mM MgCl_2_, 1 mM DTT, and 10% glycerol) to remove imidazole and stored at −80°C. The GST-His construct was manipulated under the same conditions as the negative control.

### Probe preparation for UV-crosslinking

To create the template for transcript RNA IRES_61-240_ and IRES_mut 61-240_, the pGAP1 plasmid was used as the template for PCR amplification. Two primers, T7-IRES 61+ (5′-TAATACGACTCACTATAGGGTCTACAAAAAGAAAAAAC-3′) and T7-IRES 61mut+119 (5′-TAATACGACTCACTATAGGGAGATGTTTTTCTTTTTTGGCTCATTTATATTCCTTTTCCCATTTAGTTTTGTGTCTTTTTCATCAGAAAATTCTTC-3′), along with the downstream primer EcoRI-IRES-240 (5’ <u>GAATTC</u>AAAATAGAGAGAATTGAGCTAAATTTA-3′, *Eco*RI site underlined), were used to synthesize 206 bp DNA fragments. The amplified DNA fragments were then gel purified and cloned into *Sma*I-cut pUC18. After verification, these two constructs were designated as pIRES61-240 and pIRESmut 61-240. For *in vitro* transcription, the plasmids were further digested with *Eco*RI.

### UV-crosslinking and competition assay

The purified proteins (PAB8-His or GFP-His) were incubated with labeled IRES_61-240_for 5 min at room temperature in binding buffer (10 mM Tris-HCl, pH 8.0, 150 mM NaCl, 1 mM MgCl_2_), 100 ng/μl bovine serum albumin (BSA), 50 ng/μl yeast total RNA and RNase out. The mixture was then irradiated with a 254-nm UV lamp (Stratagene, La Jolla, CA, USA, UV Stratalinker 1800) on ice for 10 min. After irradiation, the samples were treated with 2 □l RNase A (10 mg/ml) for 25 min at 37°C, boiled in 1X Laemmli buffer, and electrophoresed on a 12% SDS-PAGE gel. For competition reactions, various amounts of unlabeled competitor RNAs were pre-incubated with the proteins for 5 min before the addition of the ^32^P-labelled RNA probe.

### Protein detection and western blot analysis

Proteins were extracted from 100 mg of *N. benthamiana* leaves at 3 days post-infiltration. The extraction was carried out using 250 μl of protein extraction buffer composed of 1M Tris-HCl, pH 8.0, 3M KCl, 1M MgCl_2_, 0.5M EDTA, 60% glycerol, 10% SDS and 10% β-mercaptoethanol). The protein samples underwent a boiling water for 5 min, and subsequently, 5 μl of each sample was separated by 14% SDS-PAGE. Following separation, the proteins were transferred onto a polyvinylidene difluoride (PVDF) membrane, and subjected to western blotting analysis using the polyclonal rabbit antibodies specific to BaMV coat protein, green fluorescent protein (GFP), or actin. The PVDF membrane was then washed and incubated with secondary antibody (IRDye 800CW Gat anti-Rabbit IgG Secondary Antibody; LICORbio, Lincoln, NE, USA; Cat. No. 926-32211). Fluorescence density was detected and quantified using Amersham Typhoon (GE healthcare, Chicago, IL, USA).

### Total RNA extraction

Approximately 100 mg of *N. benthamiana* leaf tissues, collected 3 days post-infiltration, underwent cryogenic pulverization with liquid nitrogen. The resulting powder was thoroughly mixed with RNA phenol/chloroform (pH 4.3) and RNA extraction buffer (50 mM NaOAc pH 5.2, 10 mM EDTA, and 1% SDS) via vortexing. Following centrifugation at 13,200 rpm for 5 min at 25 °C, RNA extraction occurred from the aqueous layer. Subsequently, this RNA underwent a section extraction with phenol/chloroform (pH 4.3) and precipitated using 7.5 M NH_4_OAc and 100 % ethanol. The ultimate RNA pellet underwent drying, was reconstituted in 30 μl deionized water, and the resultant RNA was stored at −80 °C.

### Prepare RNA probe and northern blot analysis

To generate pGEM4-GFP/HE for northern blotting, the GFP gene was amplified via PCR utilizing the EcoRI/GFP-F (5′-GCGAATTCATGGTGAGCAAGGGC-3′) and HindIII/GFP-R (5′-GCAAGCTTTTACTTGTACAGCTC-3′) primer set. Subsequently, the resulting PCR product was cloned into the *Eco*RI and *Hin*dIII-digested pGEM-4 vector. This plasmid serves as a template for the synthesis of a ^32^P-labelled 0.7-kb RNA transcript complementary to the GFP RNA. The *in vitro* transcription reaction, conducted in a 20 μl volume, comprised 2 μg of linearized pGEM4-GFP/HE DNA with *Eco*RI, 5X transcription buffer (Promega), 10 mM DTT, 3 mM (A/C/G)TP, 250 μM UTP, T7 RNA polymerase (20 U/μl, Promega), and 70 μCi [α-32p]UTP (Dupont-NEN, Boston, MA). Total RNA extracted from *N. benthamiana* leaf tissues underwent denaturation with a glyoxal buffer (10 mM sodium phosphate buffer, pH 7.0, 50% DMSO, 1 M glyoxal), followed by a separation in a 1% agarose gel. The resolved RNA was transferred onto Zeta-probe nylon membranes (Bio-Rad Laboratories, Hercules, CA, USA; Cat. No. 162-0190) (Tsai and Dreher, 1991). For membrane probing, a □α□^32^P]UTP□labelled probe (10^6^ cpm) was employed, and subsequent hybridization was followed by scanning using an Amersham Typhoon (GE healthcare, Chicago, Illinois, USA).

### Florescence analysis

*Agrobacterium* harboring the pEpyon/mGFP-GAP5′UTR-OFP construct was co-infiltrated with FLuc/HA (control) or PAB8/HA, along with the silencing suppressor HcPro, in an 1:1:1 ratio and infiltrated into *N. benthamiana* leaves for one day. Images were captured using laser scanning confocal microscopy (Olympus Fluoview FV3000) with excitation/emission wavelength of 561 nm/586 nm for OFP. The image intensity was quantified by using the cellSens software (Olympus, Japan).

### RNA extraction and real-time PCR

When the plants reached six weeks of age, they underwent mechanically inoculation with 150 ng of BaMV on each leaf. Leaves, both virus-inoculated and mock-inoculated (with H_2_O), were harvested on days 1, 3, or 5 post-inoculation (dpi). Subsequently, the virus- and mock-inoculated *N. benthamiana* leaves were ground, and total RNA was extracted following the procedure described (Chen *et al*., 2017). The cDNA synthesis reaction adhered to the manufacturer’s instructions, utilizing ImProm-IITM Reverse Transcriptase (Promega; Cat. No. A3802), and Oligo dT25 primers.

The relative expression levels of *NbPAB8* were determined through real-time RT-PCR in a 20 µl reaction, consisting of a 1X KAPA SYBR FAST qPCR Kit (KAPA Biosystems, Boston, MA, USA; Cat. No. KK4601), and 0.2 µM of the primer pair Pab8 984 5’(+) qpcr (5′-GTTCTCTCCCTATGGCACAATC-3′) and Pab8 11653’(−) qpcr (5′-CCTGTAACCTTGCTCTTCTCTC-3′) in 0.2-ml PCR tubes. The thermal cycling conditions included an initial denaturation at 95□ for 3 min, followed by 40 cycles of 95□ for 3 sec and 60□ for 20 sec. Reactions were conducted in a TOptical Gradient 96 (Biometra, An Analytik Jena Company, Germany) with data acquisition at 60□ on the channel, excitation at 470 nm, and detection at 585 nm. A negative control, lacking template or reverse transcriptase, was included, and actin was amplified using the primer set actin_5’ (5′-GATGAAGATACTCACAGAAAGA-3′) and actin_3’ (5′-GTGGTTTCATGAATGCCAGCA-3′) for normalization.

## Results

### Identification of IRES activity within the *NbRabGAP1* 5′ UTR

The *NbRabGAP1* mRNA features an unusually long 553-nt 5′ UTR containing three short upstream open reading frames (uORFs) (Fig. 1A), suggesting that its expression may be strictly regulated. Because such complex leader sequences typically impede ribosome scanning, we hypothesized that *NbRabGAP1* might utilize an internal ribosome entry site (IRES) to facilitate translation. To test this, we employed a bicistronic reporter system (p2Luc) to evaluate translation efficiency *in vitro*.

**Fig. 1.**
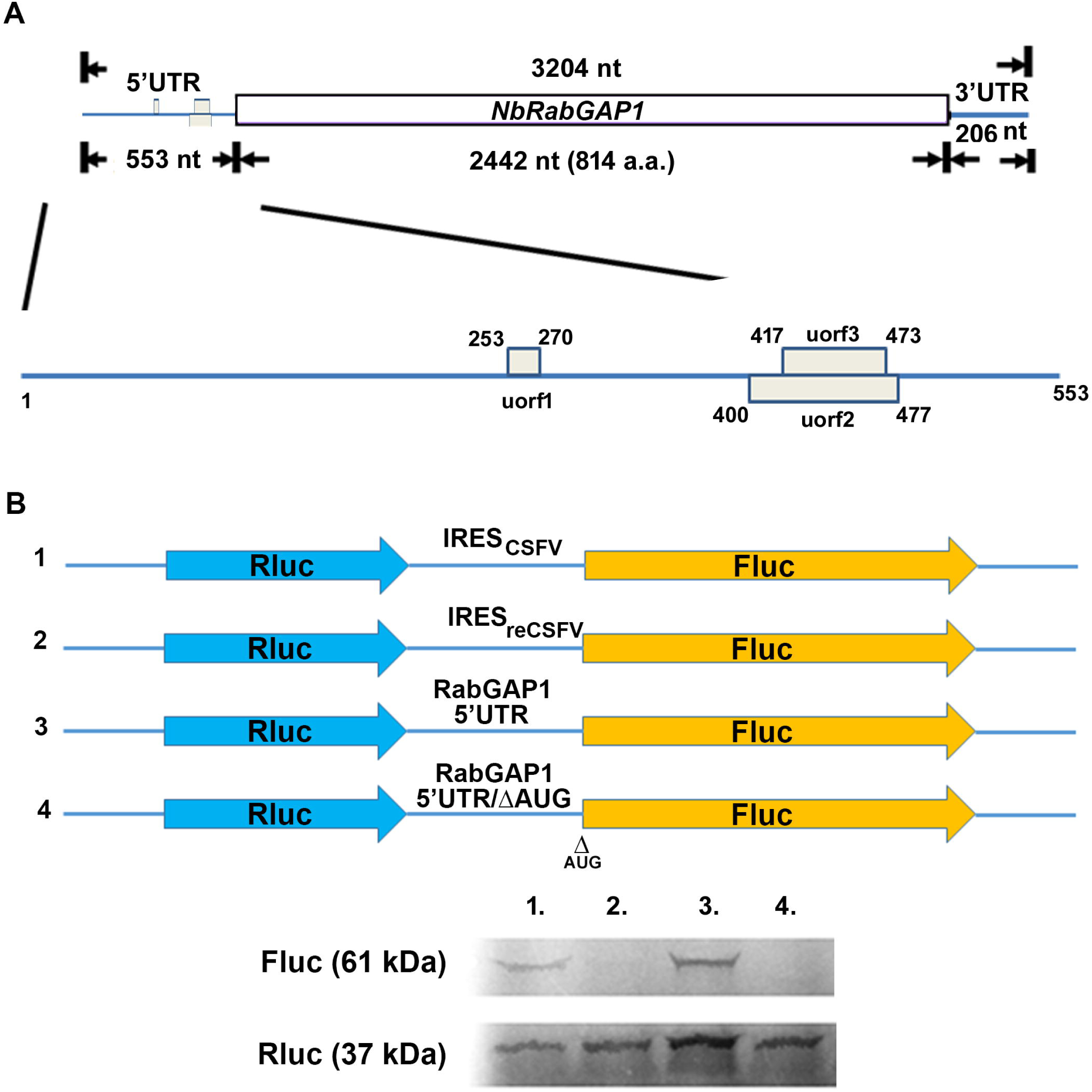
Schematic presentation of the full-length cDNA of *NbRabGAP1* and assessment of its potential internal ribosome entry site (IRES) activity using *in vitro* translation. (A) Schematic illustration of the full-length cDNA of *NbRabGAP1*. The 5′ untranslated region (UTR) is 553 nts in length and contains three upstream open reading frames (uORFs) as indicated. (B) Design of dual-luciferase bicistronic construct (p2Luc), consisting of the Renilla luciferase (Rluc) gene upstream and Firefly luciferase (Fluc) gene downstream. Construct 1 contains the intergenic region from classical swine fever virus (CSFV) IRES as a positive control. Construct 2 carries the reverse sequence of the CSFV IRES (IRES_reCSFV_) as a negative control. Construct 3 includes the full 5′ UTR of *NbRabGAP1*, and Construct 4 contains the 5′ UTR of *NbRabGAP1* with its initiation codon removed.

For initial validation, the classical swine fever virus (CSFV) IRES served as a positive control due to its established ability to drive cap-independent translation (Sizova *et al*., 1998). In our assays, the downstream firefly luciferase (Fluc) was successfully translated when driven by the CSFV IRES, whereas the reverse orientation (IRES_reCSFV_) yielded no activity (Fig. 1B).

Crucially, the *NbRabGAP1* 5¢ UTR exhibited robust IRES activity in *in vitro* translation assays. The translation was strictly dependent on the proper initiation site, as the removal of the Fluc start codon (AUG) completely abolished protein production (Fig. 1B). These findings strongly indicate that the 5′ UTR of *NbRabGAP1* contains a functional IRES that mediates internal initiation at the downstream ORF.

### Mapping the core IRES element within the *NbRabGAP1* 5′ UTR

To confirm that the translation of the downstream firefly luciferase (Fluc) reporter was driven specifically by the intercistronic region—rather than ribosome reinitiation or read-through from the upstream Renilla luciferase (Rluc) —we inserted a stable stem-loop structure (ΔG = −52 kcal/mol) immediately after the Rluc stop codon. This modification, resulting in the vector p2Luc/SL (hereafter referred to as SL) (Fig. 2A). effectively blocks scanning ribosomes. In this system, the maize *Hsp101* IRES (Dinkova *et al*., 2005; Jimenez-Gonzalez *et al*., 2014) (SL-Hsp101) served as a positive control, showing approximately 72% of the activity compared to the full-length *NbRabGAP1* 5′ UTR (SL-GAP1/1-553) (Fig. 2).

**Fig. 2.**
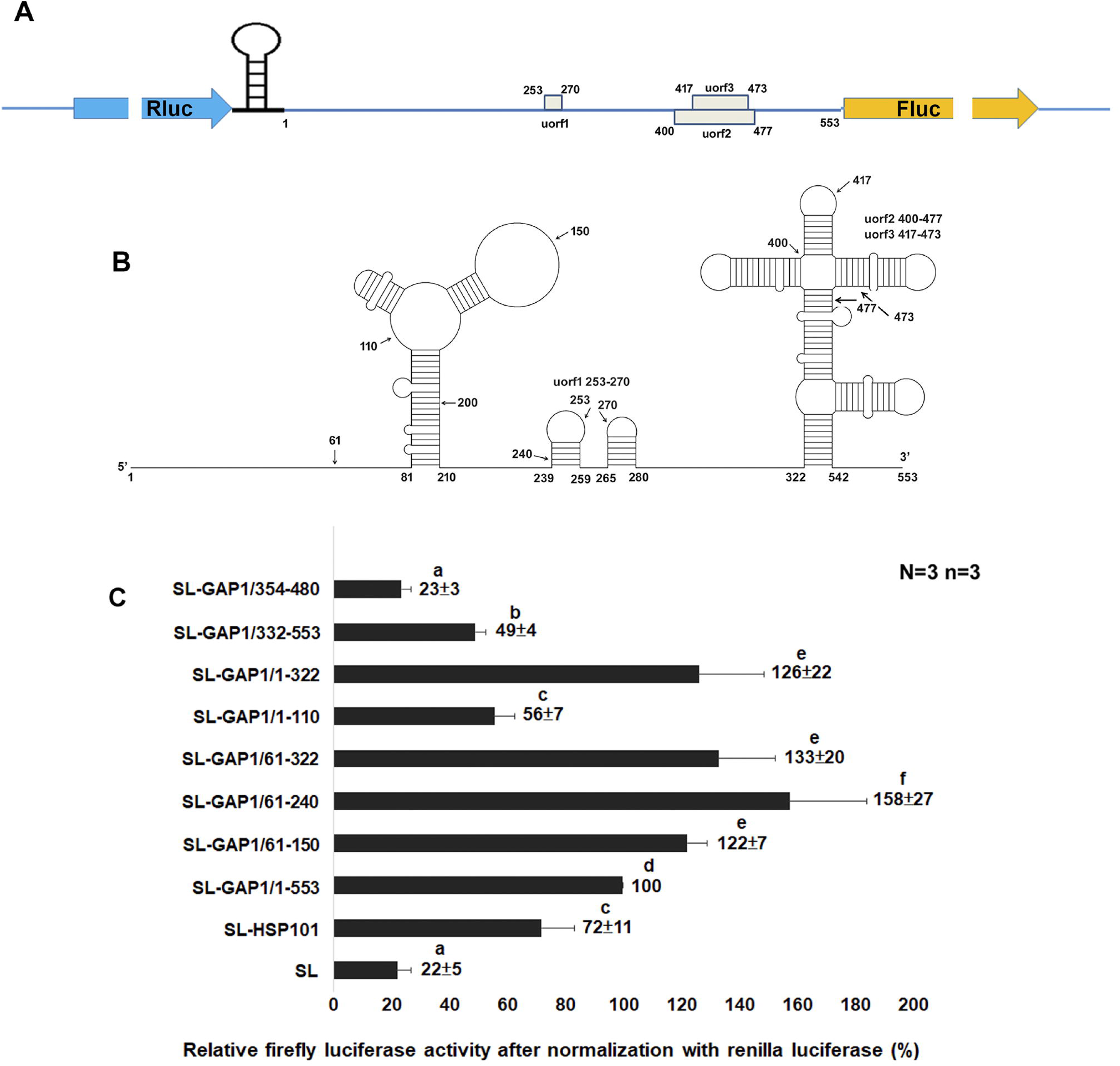
Mapping the internal ribosome entry site (IRES) activity within the 5′ untranslated region (UTR) of *NbRabGAP1*. (A) Design of the dual-luciferase bicistronic reporter (p2Luc), containing Renilla luciferase (Rluc) upstream and Firefly luciferase (Fluc) downstream, with a stable stem-loop structure inserted immediately downstream of the Rluc gene to prevent ribosome read-through. Various segments derived from the 5′ UTR of *NbRabGAP1* were inserted into the intercistronic region upstream of FLuc. (B) Predicted secondary structure of the *NbRabGAP1* 5′ UTR generated using the mfold program and visualized withe AutoCAD. (C) Relative Fluc activities (normalized to Rluc) of different *NbRabGAP1* 5′ UTR fragments: nt 354 to 480 (SL-GAP/354-480), nt 322 to 553 (SL-GAP/322-553), nt 1 to 322 (SL-GAP/1-322), nt 1 to 110 (SL-GAP/1-110), nt 61 to 322 (SL-GAP/61-322); nt 61 to 240 (SL-GAP/61-240), and nt 61 to 150 (SL-GAP/61-150). The p2Luc construct containing only the stem-loop (SL) and the construct containing a known IRES from Hsp101 (SL-Hsp101) were used as negative and positive controls, respectively. The activity of the full-length 5′ UTR (SL-GAP1/1-553) was set as 100% for comparison. Data represent means ± SE from three independent experiments (N=3), each performed in triplicate (n=3). Statistical significance was analyzed using one-way ANOVA followed by Tukey Honestly Significantly Difference (HSD) tests. Bar sharing the same lowercase letters are not significantly different (*p* < 0.05).

Using secondary structure prediction (mfold), we identified two major stem-loop regions (Fig. 2B) and divided the 553-nt UTR into specific fragments to pinpoint the core functional elements. The 5′ portion (nts 1-322) showed 126% of the full-length activity, whereas the 3′ portion (nts 322-553) retained only 49% (Fig. 2C), indicating the primary IRES activity resides in the first 322 nucleotides.

Further truncation analysis revealed that nts 61-322 were sufficient to support high translational activity (133% relative to the full-length). Among shorter fragments, nts 61-150 were identified as the minimal functional IRES, retaining 122% activity, whereas the extended fragment spanning nts 61-240 exhibited the highest translational efficiency (158%). Secondary structure prediction suggests that this region adopts a stable Y-shaped stem-loop conformation, which may facilitate efficient ribosome recruitment; however, this structural model has not yet been experimentally validated.

### BaMV infection enhances *NbRabGAP1* IRES-mediated translation *in planta*

To determine if the identified IRES functions within a living host, we employed a transient expression system in *N. benthamiana*. We compared a control construct expressing GFP alone with a GAP1/60-GFP construct, which carries the *NbRabGAP1* 5′ UTR and the first 60 nucleotides of its coding region (Fig. 3A). These were co-expressed with either an empty vector (mock) or infectious viral clones of BaMV and potato virus X (PVX).

**Fig. 3.**
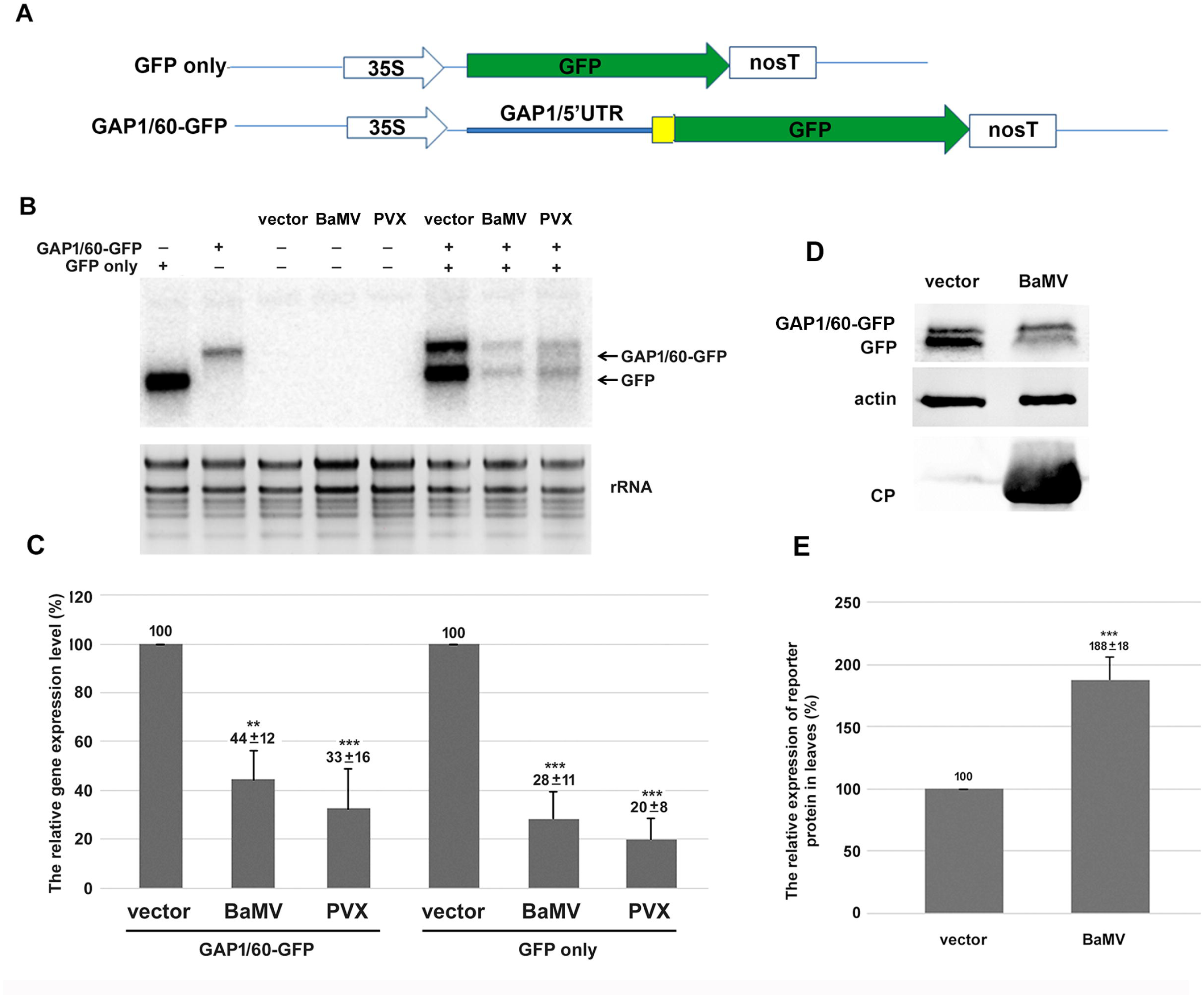
Evaluation of internal ribosome entry site (IRES) activity of *NbRabGAP1* in planta. (A) Schematic representation of reporter gene constructs: one containing the 5′ untranslated region (UTR) plus 60 nts of the NbRabGAP1 coding region fused upstream of GFP (GAP1/60-GFP), and the control construct containing GFP alone. (B) Northern blot analysis of GFP transcripts expressed in *N. benthamiana* leaves after transient expression of the constructs shown in (A), with or without bamboo mosaic virus (BaMV) or potato virus X (PVX) inoculation as indicated. (C) Quantification of relative GFP transcript levels with or without virus infection from (B). (D) Western blot analysis of GFP and GAP1/60-GFP proteins expressed in *N. benthamiana* leaves after transient expression of the constructs shown in A, with or without BaMV inoculation. (E) Quantification of relative GAP1/60-GFP/GFP protein levels with or without BaMV inoculation from (D). Data represent means ± SE from at least three independent experiments. \*\**p* < 0.01, \*\*\**p* < 0.001 by Student’s *t*-test.

Our analysis revealed several critical insights into the regulatory switch during infection. Northern blot analysis showed that transcript levels for both GFP and GAP1/60-GFP significantly decreased following BaMV or PVX infection (Fig. 3B, 3C), likely due to viral impacts on the 35S promoter. Despite the drop in mRNA levels western blot analysis revealed that GAP1/60-GFP protein levels did not decrease during BaMV infection (Fig. 3D). When normalized to the ratio of expressed GAP1/60-GFP to control GFP showed a significant twofold increase in BaMV-inoculated leaves (Fig. 3E).

These results suggest that while global cap-dependent translation may be suppressed by viral competition or host stress responses, the *NbRabGAP1* 5′ UTR actively utilizes its IRES to maintain protein synthesis. This mechanism ensures that essential host factors like NbRabGAP1 remain available to support viral movement even when the cellular translation machinery is compromised.

### NbPAB8 functions as an ITAF to enhance *NbRabGAP1* IRES activity

In Arabidopsis, certain transcripts can evade global translational repression during immune responses through R-motif-mediated translation, which recruits poly(A)-binding protein 8 (PAB8) (Wang *et al*., 2022). We investigated whether a similar mechanism facilitates the cap-independent translation of *NbRabGAP1*.

The *NbRabGAP1* IRES (nt 61-240) contains a purine-rich region where the purine content reaches ∼76% (34 out of 45) (Fig. S1). Using UV-crosslinking and competition assays with purified recombinant NbPAB8 (Fig. 4A, B), we have observed a specific interaction of NbPAB8 directly to the radiolabeled IRES_61-240_ RNA probe (nts 61-240). This binding was displaced by excess unlabeled IRES_61-240_, but not by an R-motif mutant (IRES_mut 61-240_) where purine-pyrimidine swap mutation (Fig. 4C, D).

**Fig. 4.**
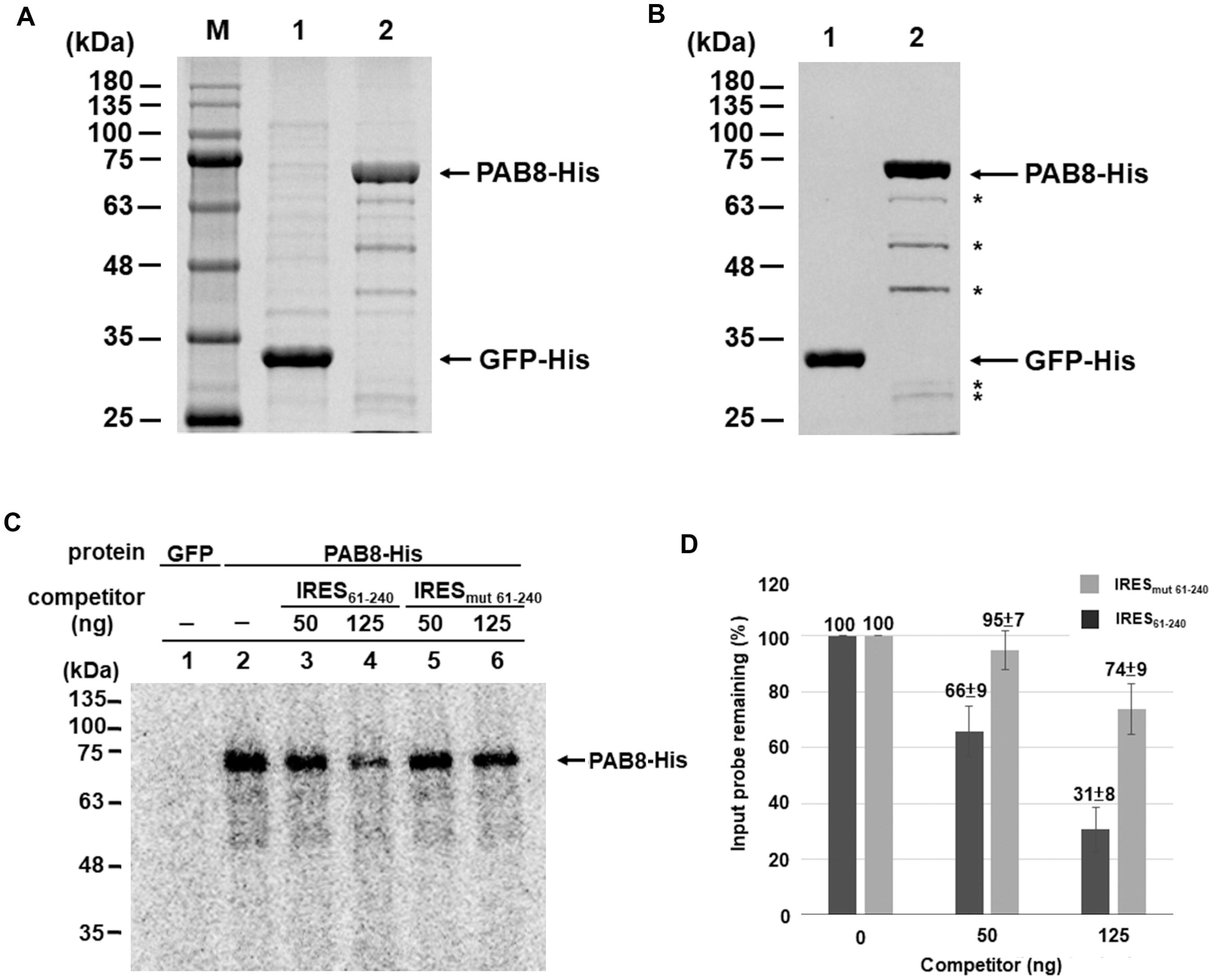
Expression of recombinant GFP-His and PAB8-His in *E. coli* and their interaction with specific RNA using UV-crosslinking assay. (A) Recombinant GFP-His and PAB8-His proteins were expressed in *E. coli* BL21and purified. The proteins were resolved on a 12% SDS-PAGE gel and visualized by Coomassie blue staining. (B) Western blotting analysis of the purified GFP-His and PAB8-His proteins using anti-His antibody. Arrowheads indicate the expected positions of GFP-His and PAB8-His. Asterisks denote degraded products derived form PAB8-His. (C) UV cross-linking assay of recombinant GFP-His or PAB8-His proteins (500 ng each) incubated with a [^32^P]-labeled RNA probe (IRES_61-240_), in the presence or absence of increasing amounts of unlabeled competitor RNAs (50 ng and 125 ng). (D) Quantification of band intensities from panel (C). The binding intensity without competitor RNA competitor was set as 100%. Data present means ± SD from at least three independent UV cross-linking experiments.

To assess functional enhancement, we utilized a bicistronic reporter expressing mGFP (upstream and Flag-tagged) and a peroxisome-target OFP (downstream and T7-tag, driven by the IRES) (Fig. 5A). Co-expression with NbPAB8-HA increased OFP protein levels by 2.8-fold compared to controls (Fig. 5B). NbPAB8 enhanced translation driven by the wild-type IRES_61-240_ by 2.6-fold, but it had no effect on the mutated R-motif (IRES_mut 61-240_) (Fig. 5C, D). Confocal imaging (Fig. 6A) confirmed that OFP fluorescence intensity within peroxisomes was ∼5.2-fold higher in the presence of NbPAB8 (Fig. 6B).

**Fig. 5.**
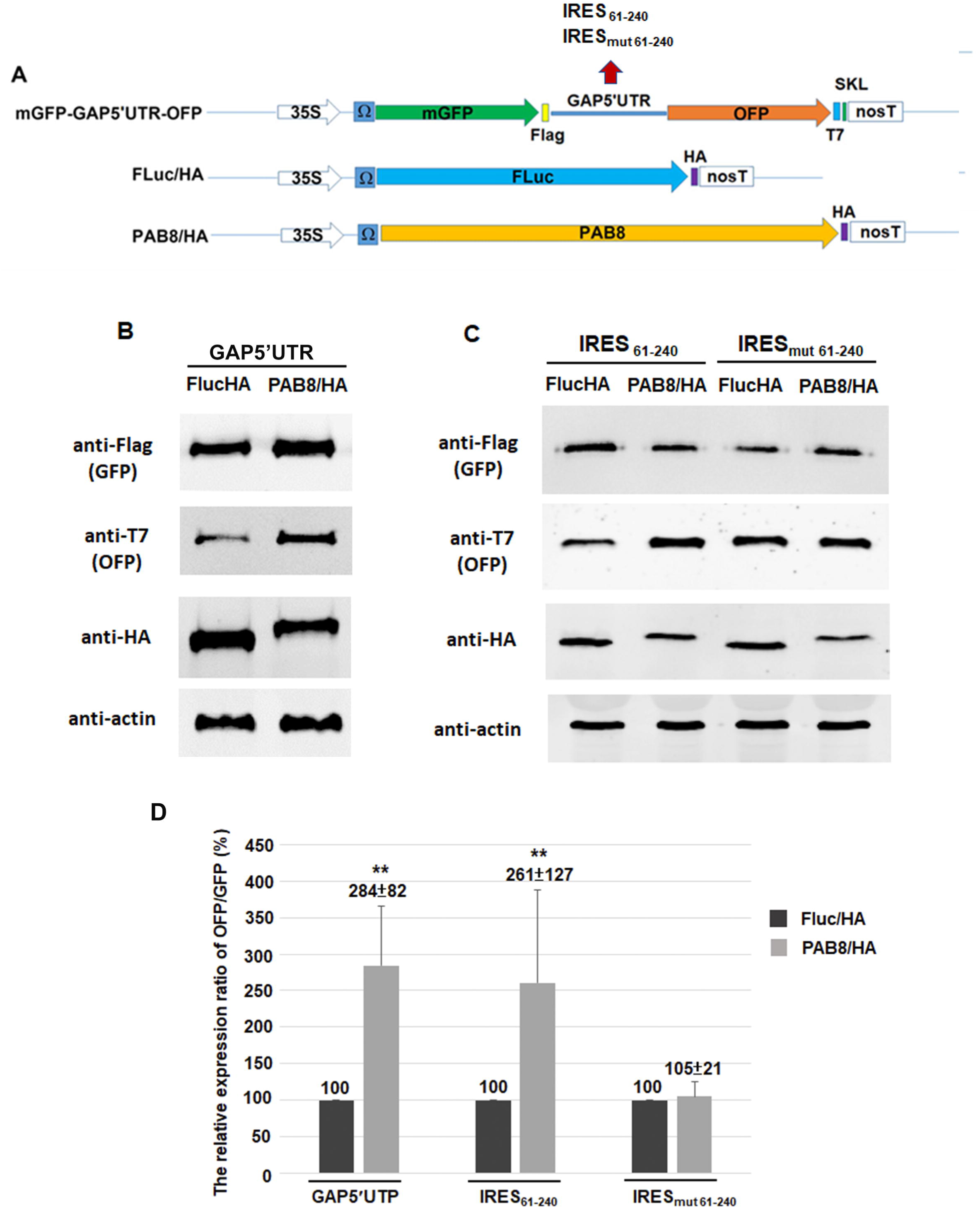
Assessment of internal ribosome entry site (IRES) activity in planta using bicistronic reporter constructs. (A) Schematic representation of reporter constructs driven by the 35S promoter and translational enhancer (Ω). The biscistronic construct consists of an upstream mGFP fused to Flag-tag, followed by either the full 5′ untranslated region (5′ UTR) of *NbRabGAP1*, the defined IRES61-240 region, or its mutant version (IRES_mut61-240_) inserted between the two cistrons, and a downstream OFP fused to a T7-tag and peroxisome-targeting signal SKL (mGFP-GAP5′UTR or IRES-OFP). Additional constructs include Firefly luciferase fused to a HA-tag (Fluc/HA), and NbPAB8 fused to a HA-Tag (PAB8/HA). (B, C) Representative western blot analysis of the expressed proteins using the following antibodies: anti-Flag for mGFP, anti-T7 for OFP, anti-HA for Fluc or PAB8, and anti-actin as a loading control. (D) Quantification of relative reporter protein expression in *N. benthamiana* leaves after co-expression with either Fluc/HA or PAB8/HA. Data represent means ± SE from at least three independent experiments. \*\**p* < 0.01 by Student’s *t*-test.

**Fig. 6.**
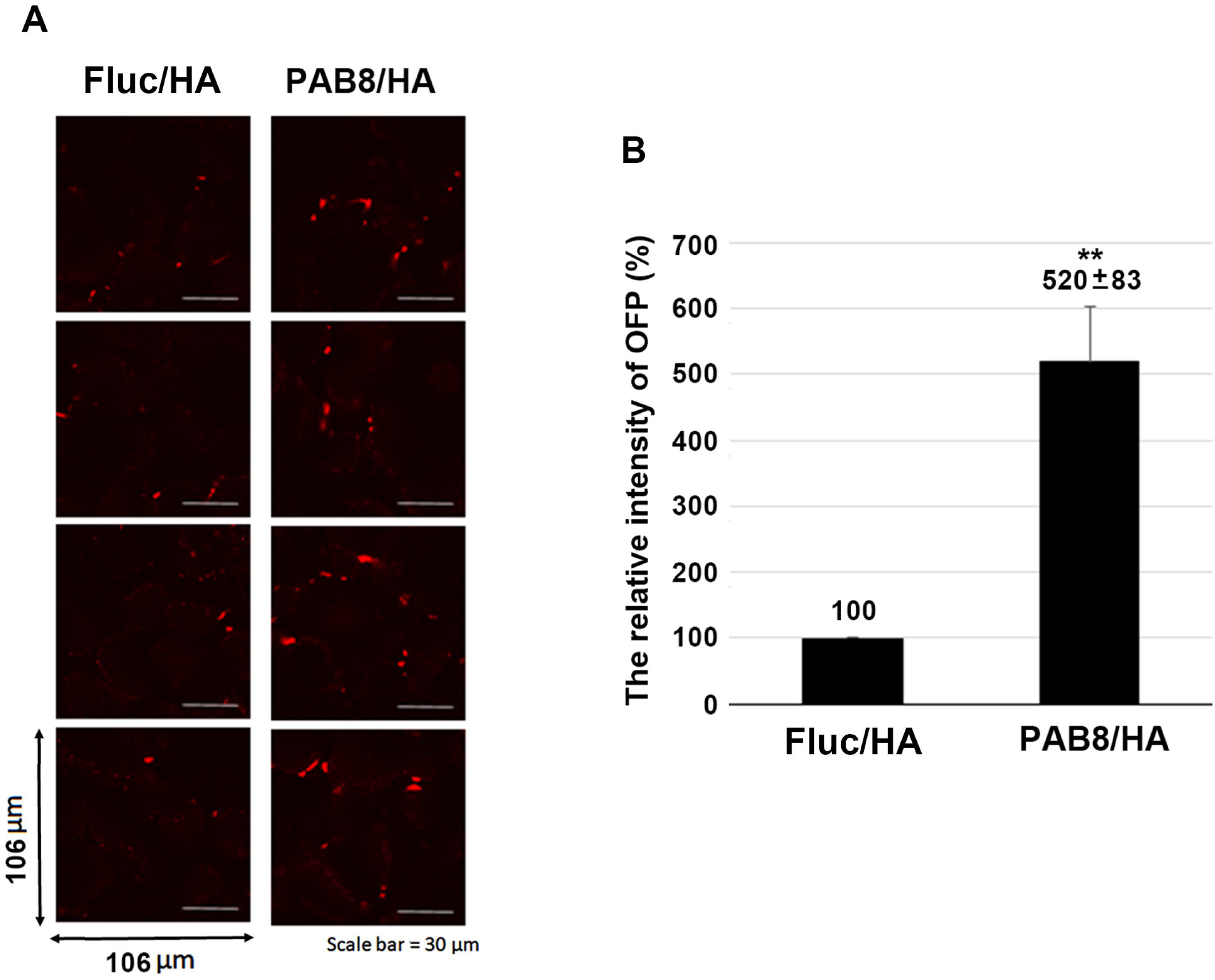
Fluorescent intensity analysis of IRES-mediated OFP expression in *N*. *benthamiana* leaves. (A) The biscistronic construct mGFP-GAP5′UTR-OFP was co-expressed with Fluc/HA or PAB8/HA in *N. benthamiana* leaves via *Agrobacterium*-mediated transient expression. The IRES-mediated OFP, translated in the cytoplasm, was targeted to peroxisomes using the SKL targeting signal. The fluorescence intensity of OFP within a 106 μm × 106 μm region was quantified from confocal images. (B) Quantification of OFP fluorescence intensity in leaves co-expressing Fluc/HA or PAB8/HA, as shown in (A). A total of 27 regions were measured for each treatment (n=27). Data represent means ± SE from at least two independent experiments (N=2). \*\**p* < 0.01 by Student’s *t*-test.

Finally, we monitored *NbPAB8* expression levels during the viral life cycle. Real-time quantitative RT-PCR revealed that BaMV infection transiently upregulates NbPAB8 transcripts, with a peak increase of nearly two-fold at 1 day post-inoculation (dpi) (Fig. 7). This upregulation ensures a sufficient supply of the IRES trans-acting factor (ITAF) to maintain NbRabGAP1 synthesis, thereby supporting efficient viral movement.

**Fig. 7.**
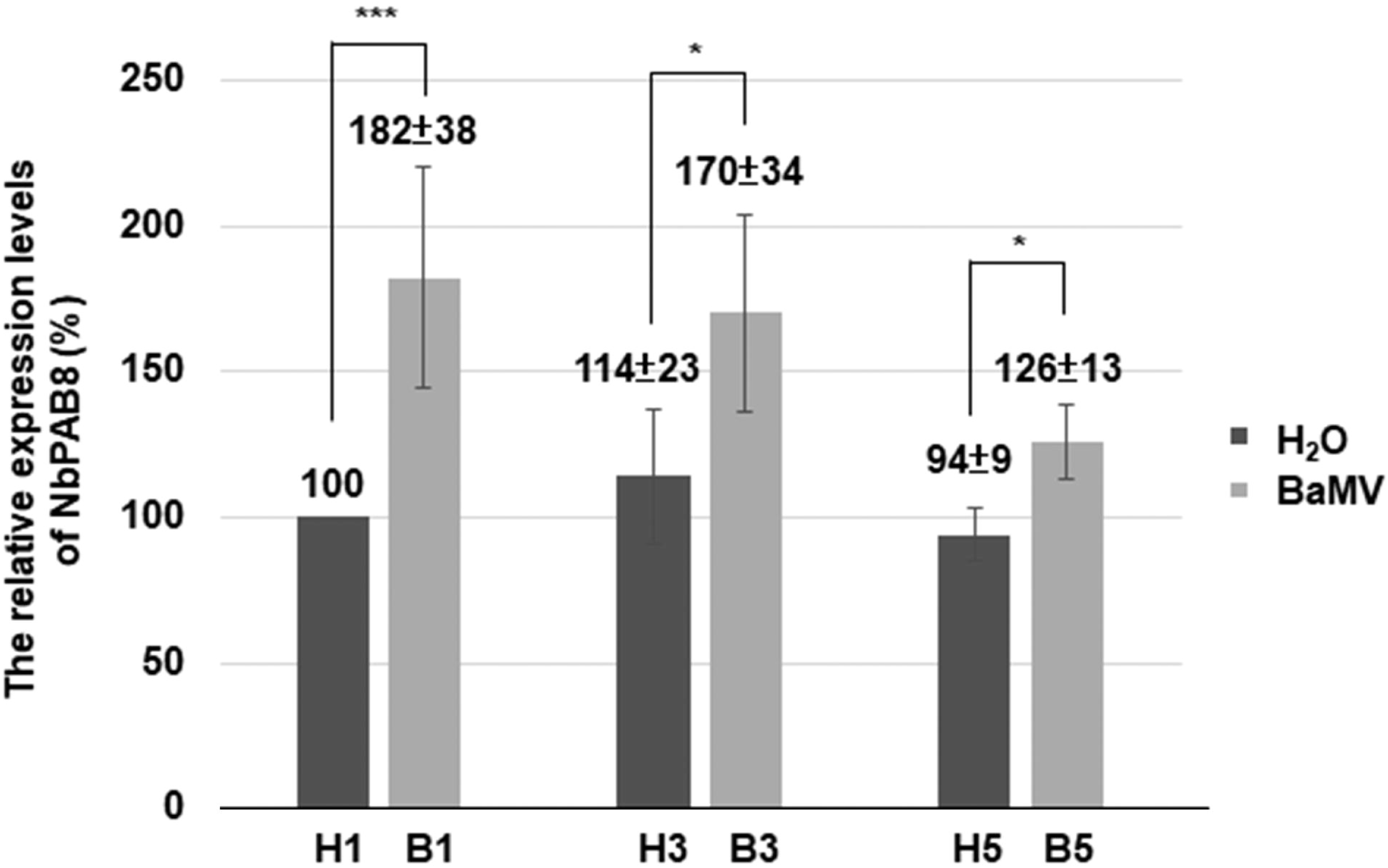
Relative mRNA expression levels of *NbPAB8* in *N. benthamiana* following BaMV inoculation. Total RNA was extracted from *N. benthamiana* leaves at 1, 3, and 5 days post-inoculation (dpi)with BaMV or mock-inoculated with water (H_2_O). Real-time RT-PCR was performed to assess the expression levels of *NbPAB8*. Data represent means ± SE from at least three independent experiments. \**p* < 0.05, *** *p* < 0.001 by Student’s *t*-test.

## Discussion

The 5′ UTR of *NbRabGAP1* is exceptionally long (553 nts) and contains three uORFs, features that typically act as major barriers to ribosome scanning and translation efficiency. Our study demonstrates that *NbRabGAP1* bypasses these inhibitory elements by utilizing a cap-independent translation mechanism. We identified a core IRES within nts 61-150, which achieves maximum efficiency when extended to include a predicted Y-shaped stem-loop structure (nts 61-240) (Fig. 2B). This architecture resembles the IRES found in the cellular immunoglobulin heavy chain binding protein (BiP) mRNA (Le and Maizel, 1997) suggesting a conserved structural strategy for internal ribosome recruitment in eukaryotes.

The 5′ region of this IRES is highly purine-rich (∼76%), acting as a specific R-motif. Our data confirms that NbPAB8—the *N. benthamiana* homolog of Arabidopsis PAB8 (Fig. S2)—binds specifically to this R-motif to boost translation over 2.5-fold. This interaction likely functions as a specialized initiation module that remains active even when the cell undergoes translational reprogramming due to stress or immune responses (Wang *et al*., 2022).

During BaMV infection, host cells typically suppress global cap-dependent translation through mechanisms such as mRNA decapping and competition for translational resources (Wang *et al*., 2022). Our findings suggest that BaMV circumvents this repression via a strategic “regulatory switch” involving the upregulation of an IRES *trans*-acting factor (ITAF). Specifically, BaMV infection transiently increases *NbPAB8* transcript levels, with a pronounce induction at 1 dpi (Fig. 7). Elevated NbPAB8 abundance likely promotes IRES-mediated translation of NbRabGAP1 under conditions where cap-dependent translation is compromised. Because NbRabGAP1 is required for NbRabF1-mediated vesicle trafficking, this mechanism enables the maintenance of intracellular transport pathways essential for efficient cell-to-cell movement of BaMV, even in the context of global host translational repression.

Our work reveals how a plant virus hijacks a host stress-response pathway—specifically the PAB8/IRES initiation module—to prioritize the production of a host factor required for its own intracellular movement.

## Conclusions

This study elucidates a specialized translational control mechanism that ensures sustained synthesis of NbRabGAP1 during BaMV infection. Although the long 5′ UTR and multiple uORFs of *NbRabGAP1* would typically impede canonical ribosome scanning, they instead harbor a sophisticated IRES element that provides an effective translational workaround under stress conditions. We identified a core IRES element located within nts 61-150, whose activity is enhanced by a Y-shaped secondary structure spanning nts 61-240. The IRES contains a purine-rich R-motif (∼76% purines) that specifically recruits NbPAB8 to initiate translation. Notably, BaMV infection transiently upregulates *NbPAB8* transcript levels, particularly at 1 dpi, thereby activating *NbRabGAP1* IRES-mediated translation. This mechanism sustains the production of vesicle trafficking machinery essential for viral cell-to-cell movement, even when global cap-dependent translation is repressed. Finally, our findings demonstrate that BaMV does not merely suppress host protein synthesis but actively rewires host translation control to selectively prioritize key host factors, such as NbRabGAP1, that are vital for its infection cycle.

## Supplementary data

**Fig. S1.** Nucleotide sequence of the NbRabGAP1 5′ untranslated region (UTR). The internal ribosome entry site (IRES) region is underlined, and purine residues within this IRES region are highlighted in yellow.

**Fig. S2.** Phylogenetic analysis of poly(A)-binding protein sequences. (A) Dendrogram depicting the evolutionary relationships among poly(A)-binding proteins from *Arabidopsis thaliana* and *N. benthamiana*, including AtPAB2, AtPAB4, and AtPAB8 from *Arabidopsis* and their homologs NbPAB2 and NbPAB8 from *N. benthamiana*. (B) Sequence similarity relationships among these poly(A)-binding proteins are highlighted. The accession numbers used for sequence retrieval are: AtPAB2 (829557 NC_003075.7), AtPAB4 (816867 NC_003071. 7), AtPAB8 (841399 NC_003070.9), NbPAB2 (Nbv6.ltrP69188), and NbPAB8 (Nbv6.1 trP68878).

## Acknowledgements

We appreciate the Bioimage Core Laboratory of the Graduate Institute of Biotechnology at National Chung Hsing University for providing the facility and the assistance. We thank Mr. Hsin-An Chen with the help of drawing the secondary structure of the 5′ UTR of *NbRabGAP1*.

## Author contributions

JLS, IHC, YWH, and YPH conceived and designed the experiments (Conceptualization, Methodology). JLS, IHC, and YPH performed experiments (Investigation). JLS, IHC, and CHT analyzed the data (Formal analysis, Data curation). YWH provided materials and technical support (Resources). JLS and IHC wrote the initial draft (Writing-original draft). All author contributed to data interpretation and manuscript revision (Writing-review & editing). CHT supervised the project and acquired funding (Supervision, Project administration, Funding acquisition).

## Data availability

All data integral to this study are presented in the figures or Supplementary datasets.

